# Early seedling development in dark conditions is directly controlled by plastids through the GUN1-dependent plastid retrograde pathway

**DOI:** 10.1101/2024.11.13.623400

**Authors:** Salek Ahmed Sajib, Michał Bykowski, Pedro Barreto, Caroline Mauve, Etienne Delannoy, Sophie Blanchet, Rim Chamas, Markus Schwarzländer, Łucja Kowalewska, Bertrand Gakière, Livia Merendino

## Abstract

In dark growth conditions, seedlings develop specific features such as an elongated hypocotyl, a tightly folded apical hook, and non-green cotyledons. This dark-specific process, known as skotomorphogenesis, relies primarily on mitochondria and eventually etioplasts for energy. Our previous research shows that skotomorphogenesis is reprogrammed in response to mitochondrial and plastidial dysfunction. Even though the direct link between mitochondria and skotomorphogenesis was described, the impact of plastid dysfunction on early development could not be separated from mitochondrial stress.

In this study, we aim to determine the direct connection between plastid functionality and skotomorphogenic response. In this situation, we analyze the phenotypic, molecular, and metabolic effects of treating etiolated seedlings using lincomycin and spectinomycin, which target plastid translation. Our results with the lincomycin treatment highlight the direct role of plastids in the control of early development, even in dark growth conditions, in the absence of any photosynthetic activity, and without the involvement of mitochondrial intermediates. Additionally, our findings suggest that GUN1 plays a regulatory role in regulating nuclear gene expression in response to plastid translation inhibition. Thanks to our study, we can now build a more precise model proposing a straight link between the reprogramming of early development and the dysfunction of plastids in dark-growth conditions.

**Significance statement:** In underground germination conditions, seedlings follow a dark-specific development program, called skotomorphogenesis, that is required for their efficient emergence from the soil. Our study demonstrates that plastids play a crucial role in controlling skotomorphogenesis, even in the absence of photosynthetic activity and without affecting mitochondrial function, and therefore we propose that regulation of etioplast functions might contribute to the adaptation of seedling development to constraining environmental conditions.

## Introduction

When the seeds are covered beneath the soil, the seedlings initiate a specific process to develop in darkness and to reach the surface toward light as quickly as possible. This developmental program, in the absence of light, is referred to as skotomorphogenesis. During skotomorphogenesis, seedlings exhibit unique characteristics, including an elongated hypocotyl, a tightly folded apical hook, and non-green cotyledons. Both intracellular and environmental signals have been shown to impact the skotomorphogenic program (Guzman and Ecker, 1990, Zhong *et al*., 2014, Abbas *et al*., 2015, Merendino *et al*., 2020, Sajib *et al*., 2023a).

During skotomorphogenesis, seed reserves are the main source of energy which are metabolized to fuel the mitochondrial respiration process, providing the required ATP for cellular processes. Etioplasts, which are small and round dark-differentiated plastids, might also supply energy in darkness through the etiorespiration process, a pathway similar to mitochondrial oxidative phosphorylation (Renato *et al*., 2015, Kambakam *et al*., 2016).

Plastids and mitochondria have prokaryotic origins and they contain their own genomes. However, they also depend on the nucleus for the expression of their proteome. Plastids contain two distinct types of transcriptional machineries: the plastid-encoded RNA polymerase (PEP) and the nucleus-encoded RNA polymerases (NEPs: RPOTp, exclusive to plastids, and RPOTmp, targeted to both plastids and mitochondria) (Pfannschmidt *et al*., 2015). The PEP is of the prokaryotic type and is targeted by the antibiotic rifampicin (Rif). Being the PEP the only prokaryotic RNA polymerase in the cell, Rif is highly specific for plastids. Alongside the PEP, the plastid translational machinery also comprises mainly components encoded by the plastid, resembling prokaryotic types (such as 70S type-ribosomes) (Marín-Navarro *et al*., 2007). Various antibiotics, including lincomycin (Lin) and spectinomycin (Spec), can limit plastid translation by specifically targeting the 50S and 30S subunits of plastid ribosomes, respectively (Ellis, 1970). Much like plastids, mitochondria possess their own genome. Still, they entirely depend on the nucleus for the transcription of their genome by the two nuclear-encoded RNA polymerases (NEPs), RPOTm, exclusive to mitochondria, and RPOTmp, also targeted to plastids (Kuhn *et al*., 2009). They do not possess any mitochondrial-encoded and prokaryotic-like RNA polymerase, and for this reason, they are resistant to antibiotics targeting transcription, such as Rif. In contrast, even though many features of the mitochondrial translational process are different from the bacterial and plastidial counterparts, mitochondrial ribosomes exhibit a structure resembling that of prokaryotes and might also be impacted by antibiotics affecting plastid translation as Lin or Spec (Hammani and Giegé, 2014).

Given their bio-energetic functions and metabolic exchanges, we previously investigated the potential engagement of both organelles in skotomorphogenesis. Our findings demonstrated that the restriction of mitochondrial and/or plastid gene expression (PGE), together with the inhibition of mitochondrial respiration, trigger a developmental response in seedlings grown in darkness (Merendino *et al*., 2020, Sajib *et al*., 2023a). This supports the involvement of both organelles in the control of skotomorphogenesis (Sajib *et al*., 2023b). In addition, when the limitation of PGE was imposed using antibiotics such as Rif, it resulted in the induction of marker genes for mitochondrial stress and in the increase of the capacity of ALTERNATIVE OXIDASE (AOX)-dependent respiration in mitochondria (Sajib *et al*., 2023a). Finally, genetic disruption of the mitochondrial-localized AOX1a enzyme was interfering with the developmental response (overbending of the apical hook) to both Rif and Spec treatments targeting distinct plastid gene-expression steps. Under this circumstance, the impact of plastid stress became inseparable from mitochondrial stress. To determine the direct link between plastid deficiency and developmental response, we have therefore searched for a plastid-specific antibiotic treatment that could induce plastid dysfunction without also provoking mitochondrial stress. In this context, we analyze the phenotypic, molecular, and metabolic effects of treating etiolated seedlings with Lin and Spec. We propose that the perception of Lin-induced plastid stress signals initiates the reprogramming of skotomorphogenesis without the involvement of mitochondrial intermediates. Our data show the specific involvement of plastids in the developmental process, even in the absence of photosynthetic activity.

## Results

### The restriction of plastid translation by Lin and Spec treatments alters etioplast gene-expression and ultrastructure, and seedling skotomorphogenesis

To investigate the impact of PGE limitation on the morphogenesis of dark-grown seedlings, plastid translation was inhibited by treatment with specific antibiotics, namely Lin and Spec. In both antibiotic-treated seedlings, plastid-encoded RBCL (large subunit of RUBISCO) protein levels were undetectable (Figure 1 (a)). Consistently, inhibition of plastid translation resulted in a decrease of plastid transcripts synthesized by the plastid-encoded RNA polymerase (PEP), such as *RbcL* and *PsbA* (Figures 1(b)). In contrast, the levels of NEP-encoded plastid transcripts, *RpoB* and *Rpl20,* were increased as a compensation effect or not affected as for *Rpl20* after Spec treatment. We also examined the impact of PGE inhibition by both antibiotics on the plastid ultrastructure in the cotyledons of etiolated seedlings (Figure 1 (c) and Suppl. Figure 1). Untreated seedlings displayed a well-developed and highly regular bicontinuous prolamellar body (PLB) structure. However, in seedlings treated with both antibiotics, the PLB structure was disrupted, and membranes formed irregular sponge-like spots with numerous plastoglobules. The antibiotics-treated seedlings also showed a significant accumulation of starch and a decrease in the Pchlide levels, which correlates with the irregular PLB formation in the etioplast in those PGE-limited conditions (Suppl. Figure 1 (e)-(f)). Moreover, we also checked the etioplast ultrastructure in the seedlings treated with Rif, which limits plastid transcription by specifically inhibiting the activity of the PEP. In Rif-treated seedlings, no PLB formation was detected, and the amount of cytoplasm-localized lipid droplets stored in this sample was high (Suppl. Figure 2).

**Figure 1.**
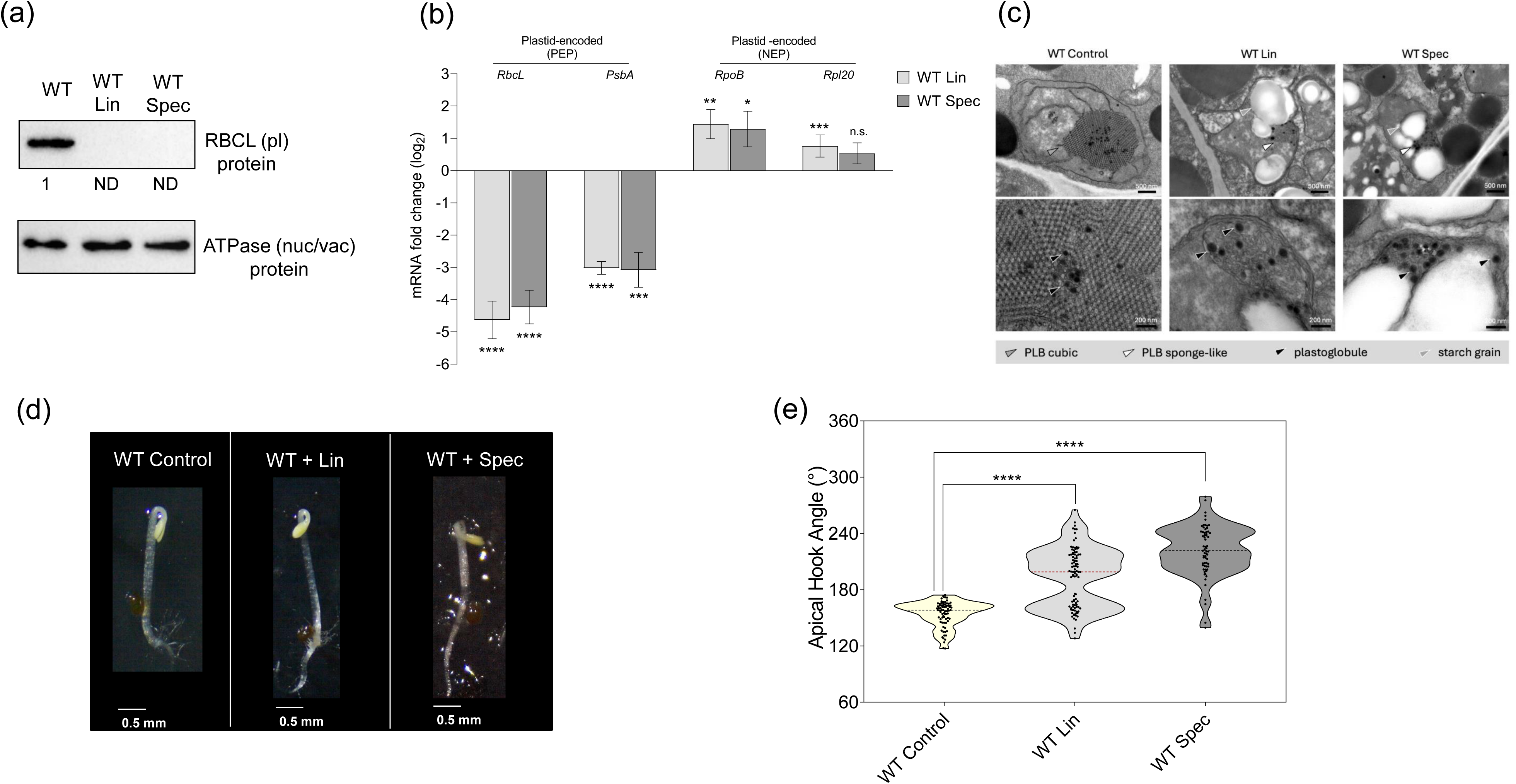
Inhibition of plastid translation by Lin and Spec impairs plastid biogenesis and reprograms skotomorphogenesis in 3-day-old etiolated Arabidopsis WT seedlings. (a) Analysis of plastid-encoded RBCL protein content in etiolated WT seedlings grown in the absence or presence of Lin or Spec: Protein extracts (20 µg) were separated by SDS-PAGE and immunoblotted with specific antisera against plastid-encoded RBCL proteins and the vacuolar ATPase proteins (nucleus-encoded), that were used as a loading control. The RBCL signals were quantified using Image Lab software with ATPase for normalization and results are shown below the RBCL protein signals. ND indicates not detected. (b) Expression analysis of plastid genes that are transcribed by PEP (Plastid Encoded RNA Polymerase) and NEP (Nuclear Encoded RNA Polymerase): Log2 fold change (FC) values of expression levels in Lin/Spec-treated WT seedlings compared to untreated seedlings were determined by reverse transcription-quantitative PCR. Expression levels were normalized to the mean (nuclear) expression of PP2A, used as a reference gene. The mean values of three biological replicates are plotted with error bars representing standard deviation. A t-test determined the statistical significance of differences (*P < 0.05; **P < 0.01, *** P < 0.001, ****P < 0.0001, n.s. = not significant). (c) Ultrastructure of etioplasts in cotyledon regions of WT seedlings grown with or without Lin/Spec. Grey arrowhead with black outline indicates the cubic prolamellar body (PLB), white arrowhead indicates PLB sponge-like structures, black arrowhead indicates plastoglobules and grey arrowhead with white outline indicates starch grains. Top panel scale bar is 500 nm; bottom panel scale bar is 200 nm. (d) Dissection microscope images of etiolated WT seedlings grown with or without Lin/Spec (the scale bar represents 0.5 mm) and (e) corresponding violin plots of median apical hook angle values: The number of pooled individuals (n, measured in two independent analyses) includes 76 seedlings for WT control, 95 for WT + Lin, and 65 for WT + Spec. All differences were statistically significant (Pearson’s Chi-squared test with Yates’ continuity correction, \*\*\*\**P* < 1.433e-07).

Inhibition of plastid translation and biogenesis resulted in a significant alteration of skotomorphogenesis (Figure 1 (d)), particularly evident in the over-bending of the apical hook (apical hook angle greater than 180°, twisting phenotype) (Figure 1 (e)). Taken together, these data support the link between the alteration of plastid gene expression/biogenesis and reprogramming of the developmental response, highlighting the role of plastids in the control of skotomorphogenesis.

### Mitochondrial biology is altered by Spec treatment but not by Lin

To determine whether Lin and Spec treatments also have an impact on mitochondrial translation, we have analyzed the levels of mitochondrial-encoded NAD9 protein by western blot. In Lin-treated seedlings, NAD9 levels were slightly increased to 125% when normalized to ATPase amounts, while Spec treatment reduced NAD9 levels to 61% (Figure 2 (a)). These findings indicate that Spec has an inhibitory effect on mitochondrial translation, while Lin exclusively restricts plastid but not mitochondrial translation. In this context, we also analyzed the plastid specificity of Rif, and as for Lin, we did not detect any inhibitory impact on the levels of the mitochondrial-encoded proteins NAD9 (Suppl. Figure 3).

**Figure 2.**
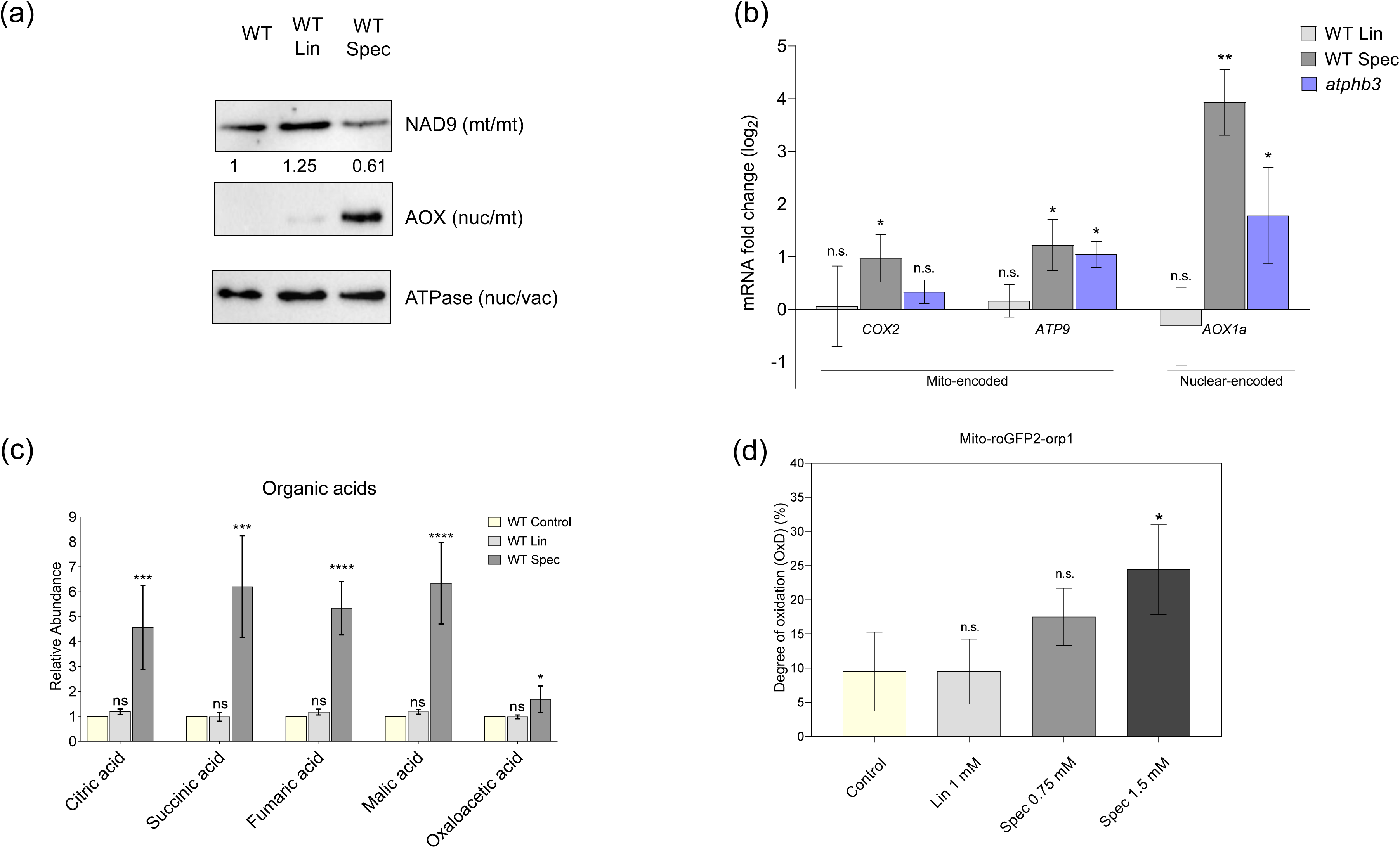
Effect of plastid translation inhibition by Lin or Spec on mitochondrial biology in 3-day-old etiolated Arabidopsis seedlings. (a) Analysis of mitochondrial protein content: Protein extracts (20 µg) were separated by SDS-PAGE and immunoblotted with specific antisera against mitochondrial-encoded mitochondrial NAD9, nuclear-encoded mitochondrial AOX and nuclear-encoded vacuolar ATPase proteins as a loading control. Quantification of the NAD9 was performed using Image Lab software with ATPase for normalization, and values are shown below the NAD9 protein signals. (b) Analysis of mitochondrial-encoded and nuclear-encoded transcripts for mitochondrial proteins. Log2(FC) values of expression levels in Lin/Spec-treated WT seedlings and untreated mitochondrial mutant *atphb3 versus* untreated WT seedlings were determined by reverse transcription-quantitative PCR. Expression levels were normalized to the mean (nuclear) expression of *PP2A*, used as a reference gene. The mean values of three biological replicates are plotted with error bars representing standard deviation. A t-test determined the statistical significance of differences (*P < 0.05; **P < 0.01, n.s. = not significant). (c) The relative abundance of organic acids was determined by GC-MS analysis in untreated and Lin/Spec-treated WT seedlings with four biological replicates. Error bars represent standard deviation. Statistically significant differences were determined using one-way ANOVA with Tukey post hoc test (n = 4; *P < 0.05, ***P < 0.001, ****P < 0.0001, n.s. = not significant). (d) Measurements of H_2_O_2_ and thiol redox status in the mitochondrial matrix: Using the mito-roGFP2-Orp1 line in control, 1 mM Lin, 0.75 mM Spec- and 1.5 mM Spec-treated conditions. The degree of oxidation was calculated from measured ratios following in situ calibration with 50 mM H_2_O_2_ and 100 mM DTT (n = 80; significantly different with *P < 0.05, by two-tailed, unpaired Student’s t-test).

Moreover, treatment with Spec, but not Lin, results in the accumulation of AOX proteins, a classical mitochondrial stress marker (Figure 2 (a)). Nuclear-encoded marker transcripts for mitochondrial stress such as *AOX1a*, *AT12CYS,* and *NDB4* in Spec-treated seedlings, but not in Lin-treated, exhibit a pattern consistent with that of the *atphb3* (*Arabidopsis thaliana Prohibitin 3*) mutant, a mitochondrial respiration mutant (Van Aken *et al*., 2016) (Figure 2 (b) and Suppl. Figure 4). These data indicate the induction of mitochondrial stress by Spec but not by Lin treatment. Additionally, levels of transcripts encoded by mitochondria (*COX2*, *ATP9*) are slightly induced by Spec treatment, similar to what was observed in the mitochondrial *atphb3* mutant, likely as a compensatory response to mitochondrial stress. At the same time, they remained unchanged when seedlings were treated with Lin (Figure 2 (b)).

We then investigated if PGE-limitation by Lin and Spec treatments has an impact on mitochondrial metabolism. We determined the changes in cellular metabolic compounds with particular attention to mitochondrial metabolites, employing gas chromatography-mass spectrometry (GC-MS). The metabolic analysis disclosed an increase in various tricarboxylic acid (TCA) intermediates, including citrate, succinate, fumarate, malate, oxaloacetate, and also glycolysis intermediates such as glucose-6-posphate and fructose-6-posphate in Spec-but not in Lin-treated seedlings (Figure 2 (c), Supplementary Figure 5 (a), table S2). Previous research also showed alterations in some of the TCA metabolites in situations of mitochondrial dysfunction, as observed in the cases of *rpoTmp* and *atphb3* mutants (Van Aken *et al*., 2016). In addition, lactic acid shows a tendency to accumulate in Spec-but not in Lin-treated seedlings (Suppl. Figure 5 (b)), suggesting the inhibition of aerobic respiration metabolism and a consequent shift to fermentation. Finally, we observe the accumulation of various fatty acids, e.g., palmitic acid and myristic acid, in Spec-treated seedlings but not in those treated with Lin (Suppl. Figure 5 (c)). All these data suggest that Spec treatment alters mitochondrial respiration metabolism, either directly or indirectly.

Increase of pyruvate due to alteration in mitochondrial metabolism might also lead to accumulation of free amino acids. To quantify these nitrogen metabolites, we employed o-phthaldialdehyde (OPA) derivatization followed by high-performance liquid chromatography (HPLC). Previously, we observed that Rif treatment alters amino acid metabolism, likely as a consequence of mitochondrial dysfunction (Sajib et al., 2023). In this study, Spec-treated seedlings exhibited a similar pattern to Rif treatment, showing significant alterations in amino acid metabolism (see Supplementary Figures 5 (a)). Pyruvate-derived branched-chain amino acids, including alanine (ALA), leucine (LEU), valine (VAL), serine (SER), and tryptophan (TRP), were accumulated in Spec-treated seedlings, likely reflecting the accumulation of glycolytic intermediates and TCA derivatives. On the other hand, there was a decrease in the levels of asparagine (ASN) and glutamine (GLN), likely due to insufficient ATP from mitochondrial respiration required for their synthesis. No relevant changes in the levels of these amino acid were observed with Lin treatment. Alternatively, aspartate (ASP) is primarily used in the plastid to support de novo synthesis of other amino acids such as homoserine (HSER), lysine (LYS), methionine (MET), and threonine (THR). In our study, we observed reduced levels of asparagine (ASN) and an accumulation of isoleucine (ILE) in Lin and Spec treatments. These findings could indicate dysfunction within the plastid in both treatments.

To evaluate the potential involvement of mitochondrial signals in the response to PGE limitation, we analyzed the mitochondrial oxidative status in seedlings expressing the H_2_O_2_ biosensor roGFP2-Orp1 in the mitochondrial matrix in the presence of Lin, Spec, and Rif. We quantified the 400/482 nm fluorescence ratio for 3 hours, and calibrated the ratio data to calculate the degree of oxidation (OxD) of the biosensor (Schwarzländer *et al*., 2008). OxD is significantly increased in the Rif-treated seedlings as also supported by the relative increase in the ratio of the signals at 400 nm over 482 nm (Supplementary Figures 6(a) and (b)). A trend towards an increase is also observed for OxD in Spec-treated seedlings (Spec 0.75 mM, Figure 2(d) and Supplementary Figure 6(c)). However, a significant augmentation in OxD and the ratio 400/482 was detected when seedlings were treated with higher Spec concentration (1.5mM). Conversely, Lin treatment showed no significant effect on mito-roGFP2-Orp1 OxD and 400/482 ratio. Our findings suggest a differential impact of these antibiotics on mitochondrial redox balance, with Spec (high concentration) and Rif inducing mitochondrial oxidative stress, while Lin appears not to alter mitochondrial redox status. These data together indicate that Lin acts on plastid gene function without significantly altering mitochondrial biology.

### The response to Lin and Spec-mediated PGE inhibition is mediated by the GUN1-dependent organellar retrograde pathway at the nuclear expression and developmental levels

Treatment with both antibiotics results in the downregulation of Photosynthetic-Associated Nuclear genes (PhANGs) such as *LHCB1*.*2* and *GLK1* in WT seedlings, as revealed by RT-qPCR analyses (Figure 3 (a)), indicating that the plastidial retrograde pathway is active even in dark conditions, as previously reported (Martin *et al*., 2016, Hernández-Verdeja *et al*., 2022). For that, we investigated the potential involvement of the plastid retrograde signaling pathway GUN1, which is responsible for the communication between the plastid and nucleus in the PGE deficiency condition (Hernández-Verdeja *et al*., 2022). The genetic impairment of the GUN1-dependent retrograde signaling pathway abolishes the Lin-dependent downregulation of the PhANGs (Figures 3(a)). In addition, genetic disruption of GUN1 also strongly reduced the size of the class including twisted seedlings (Figures 3 (b) and (c)). Intriguingly, in *gun1* seedlings, *LHCB1.2* and *GLK1* gene expression was still downregulated by Spec treatment and displayed a developmental response (i.e., over-bending of the apical hook). However, the distribution of population in the twisted and untwisted classes was inversed in Spec-treated *gun1* seedlings when compared to treated WT seedlings. These results indicate that a GUN1-dependent signaling pathway is involved in both Lin and Spec treatment, as well as a GUN1-independent pathway in the response to Spec.

**Figure 3.**
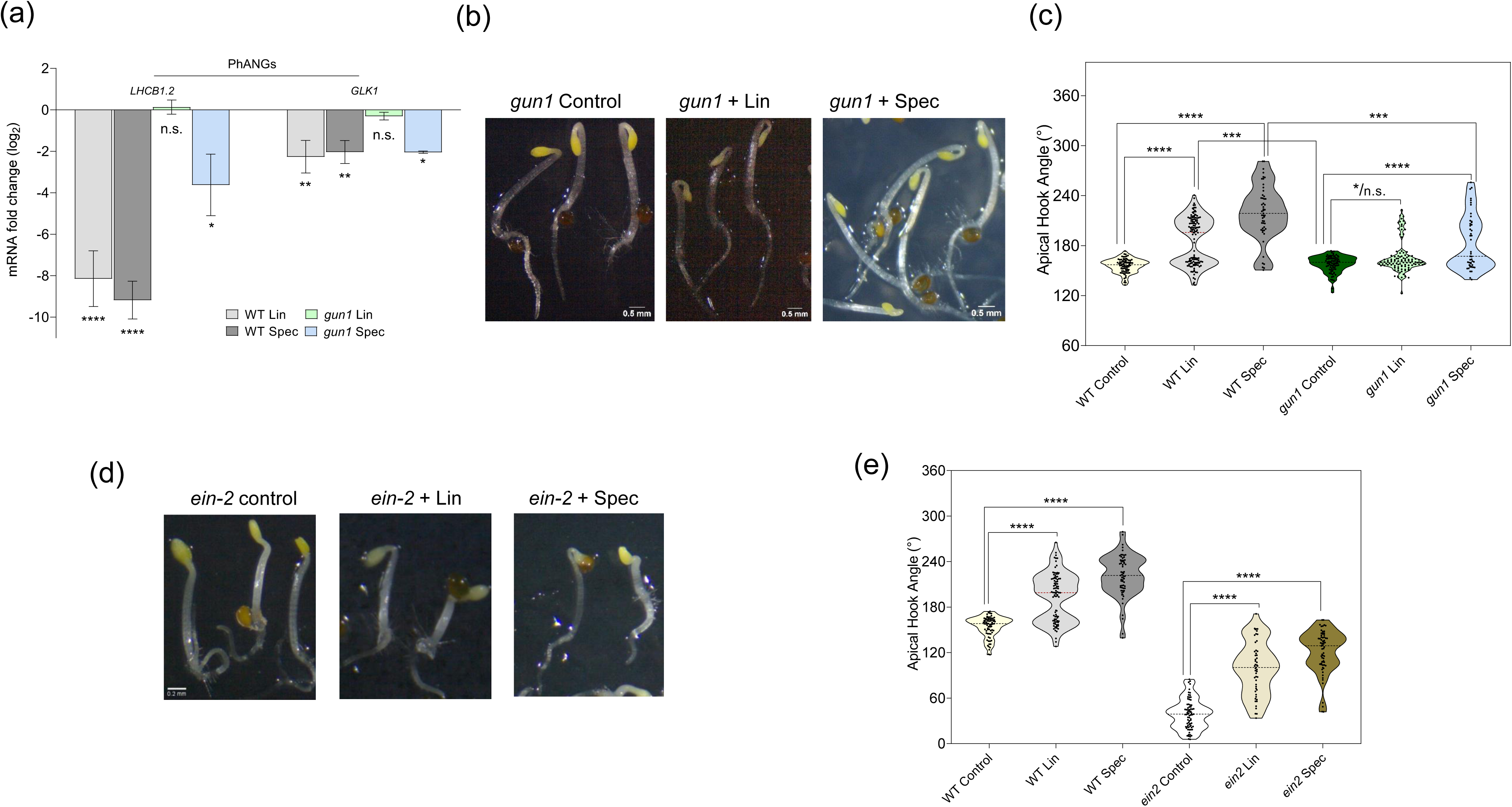
The response to plastid translation inhibition by Lin/Spec at the nuclear expression and developmental levels depends on the GUN1-mediated organelle retrograde pathway and does not require EIN2-mediated signalling pathway. (a) Expression levels of Photosynthesis-Associated Nuclear Genes (PhANGs): Log2 (FC) values in Lin/Spec-treated WT (Col-3) and mutant *gun1* seedlings versus corresponding untreated seedlings were determined by reverse transcription-quantitative PCR. Expression levels were normalized to the mean expression of PP2A, used as a reference gene. The mean values of three biological replicates are plotted with error bars representing standard deviation. The t-test determined the statistical significance of differences (*P < 0.05; **P < 0.01, ****P < 0.0001, n.s. = not significant). (b) Dissection microscope images: etiolated *gun1* seedlings were grown under control conditions, with Lin and with Spec. Scale bar corresponds to 0.5 mm. (c) Corresponding violin plots of median apical hook angle values: etiolated WT (Col-3) and *gun1* seedlings were analyzed. The number of pooled individuals (n) measured in two independent replications, which includes 80 seedlings for WT control, 132 for WT + Lin, 51 for WT + Spec, 82 for *gun1* control, 106 for *gun1* + Lin, and 46 for *gun1* + Spec. All differences were statistically significant for WT (Col-3); the *gun1* mutant was insensitive to 1 mM Lin for one replication (n.s.) but impacted by Lin in another replication (*P < 0.05). (c) Dissection microscope images of etiolated *ein2* seedlings grown with or without Lin/Spec (the scale bar represents 0.2 mm), and (e) Violin plot of median apical hook angle values. The number of pooled individuals (n) measured in two independent hook angle analyses includes 77 seedlings for *ein2* control, 51 for *ein2* + Lin, and 66 for *ein2* + Spec. Significant differences between treated and untreated seedlings were identified using Pearson’s Chi-squared test with Yates’ continuity correction (****P < 0.0001, n.s. = not significant).

### Developmental responses induced by Lin and Spec are not dependent on the EIN2-mediated ethylene signaling pathway

Earlier, we did not find any involvement of EIN2-mediated ethylene signaling pathways in the Rif-induced developmental response (Sajib *et al*., 2023a). To determine if the Lin/Spec-induced developmental response depends on EIN2-signaling pathway, we examined the skotomorphogenic profile of ethylene-insensitive mutant *ein2*-*1* seedlings exposed to antibiotics. Even though, upon treatment, these seedlings do not display the characteristic twisting phenotype when grown in the dark, they can form the apical hook, a feature that is absent in control seedlings (Figures 3 (d) and (e)). Furthermore, the downregulation of Photosynthesis-Associated Nuclear Genes (PhANGs) in PGE-limited *ein2-1* seedlings provides evidence that also the nuclear response to PGE limitation is independent of an EIN2 signaling pathway (Suppl. Figure 7).

## Discussion

Mitochondria and plastids are bioenergetic organelles that supply energy to cells for essential processes and are highly interconnected together with the rest of the cell. These semi-autonomous organelles contain their own genome and gene-expression machinery but still depend on the nucleus for the expression of most of their proteins. Retrograde signaling pathways ensure the communication between each organelle and the nucleus, allowing organelles to control the nuclear expression of their proteome according to their functional status (Pfannschmidt *et al*., 2020). Retrograde pathways can also allow the crosstalk among plastids and mitochondria with the intermediate of the nucleus (He *et al*., 2023).

It was previously found that mitochondrial dysfunction at the level of gene expression and/or respiration efficiency induces alteration of seedling architecture during skotomorphogenesis (Wang and Auwerx, 2017, Merendino *et al*., 2020). Limitation of PGE was also found to reprogram the developmental profile in darkness (Sajib *et al*., 2023a). However, the impact of plastid dysfunction on skotomorphogenesis could not be separated from mitochondrial stress. In these circumstances, it is hard to functionally link the developmental reprogramming to either plastid or mitochondrial stress. We have, therefore, searched for conditions where plastid functionality is affected without altering mitochondrial activities.

Mutations impacting essential plastidial functions for the cell can be lethal either at the embryonic or the seedling level. Mutants in the plastid-encoded RNA polymerase-associated proteins (PAPs) display a genetic arrest in chloroplast biogenesis, leading to an albino phenotype in the light (Grübler *et al*., 2017). However, in dark conditions, phenotypic observation cannot allow a distinction between mutant seedlings and WT. Alternatively, mutations can also target redundancies but might only have a weak impact on plant physiology. In this scenario, it is convenient to use antibiotic treatments to limit the prokaryotic-like PGE. Antibiotic action is directed to all seedlings, and the inhibitory effect can be modulated by adapting the dose. Antibiotics that are currently used to limit PGE target either PEP-dependent transcription or translation of the plastid transcripts. In our previous studies, we have already tested the Rif antibiotic that blocks PEP, but mitochondrial biology was found to be altered (Sajib *et al*., 2023a). Rif is plastid-specific since it targets the PEP, which is the only prokaryotic-like RNA polymerase in the cell. However, the impact of Rif is restricted exclusively to PEP-transcribed mRNAs, and NEPs can still synthesize plastid transcripts.

Conversely, the effect of antibiotics inhibiting plastid translation is broader as they target all the plastid transcripts. However, the translational machinery of mitochondria is also of the prokaryotic type, and antibiotics targeting plastid ribosomes might also affect mitochondrial translation (Hammani and Giegé, 2014). Controls of specificity are therefore required.

Lincomycin (Lin) and spectinomycin (Spec) inhibit distinct targets within the translational machinery. Lin disrupts the elongation of short peptide chains by inhibiting peptidyl transferase, while Spec impairs the translocation of peptidyl-tRNA (Burns and Cundliffe, 1973, Spahn and Prescott, 1996). Lin has long been employed to investigate nuclear responses to plastid gene expression (PGE) inhibition and retrograde signaling mechanisms (Sullivan and Gray, 1999). Previous studies have indicated that Lin does not affect translation or mitochondrial biogenesis, whereas spectinomycin may have a minor influence on cytosolic translation (Sullivan and Gray, 1999, Mulo *et al*., 2003, Doyle *et al*., 2010). Conversely, our earlier research demonstrated that Spec significantly inhibits plastid translation in 3-day-old etiolated seedlings and has also a slight impact on mitochondrial protein accumulation (Sajib *et al*., 2023a). However, a complete understanding of the effects of both antibiotics on organellar biology remains to be obtained. In this study, we investigated the effect of Lin and Spec on plastidial, mitochondrial, and nuclear gene expression, as well as intracellular ultrastructure, redox status, metabolism, and, ultimately, developmental response in the dark.

In our study, we show that treatments with either Lin or Spec block plastid translation in etiolated seedlings, as indicated by the total absence of signals in the western blot analysis of the plastid-encoded rubisco large subunit, RBCL protein (Figure 1 (a)). As a consequence of the synthesis inhibition of the plastid-encoded proteins, plastid transcription by the plastid-encoded RNA polymerase PEP is also blocked but not by the nucleus-encoded RNA polymerases NEPs, as indicated by the RT-qPCR data (Figure 1 (b)). As a result of the PGE blockage, the expression of PhANGs was downregulated in both treatments, which is an indication of plastid stress (Figure 3 (a)) (Sullivan and Gray, 1999). In addition, the ultrastructural features of the PLB were disrupted, which suggests the interference of all tested antibiotics (Spec, Lin, Rif) with plastid biogenesis in seedlings (Figure 1C, Supplementary Figures 1 and 2). Previous studies have shown that the treatment of norflurazon, lincomycin, and spectinomycin causes notable structural changes in both etioplasts and proplastids (Doyle *et al*., 2010, Martin *et al*., 2016). Moreover, the proplastids treated with norflurazon showed plastoglobuli with irregular shapes (Doyle *et al*., 2010). We also found a similar type of response where treatment with lincomycin and spectinomycin (Lin/Spec) resulted in the formation of larger plastoglobuli and the accumulation of starch granules (Supplementary Figures 1). Protochlorophyllide (Pchlide) is the most abundant among different types of PLB pigments (Bykowski *et al*., 2020). The Pchlide:LPOR: NADPH complex is known to be associated with the PLB structure formation (Dehesh and Ryberg, 1985, Böddi *et al*., 1989), and plants presenting a decrease in Pchlide accumulation do not develop PLBs (von Wettstein, 1991, Franck *et al*., 2000). Consistently, in our study, we found that the Pchlide amount is significantly decreased in the Lin/Spec treated seedlings (Supplementary Figures 1 (e) and (f)).

We then verified the impact of the antibiotic treatments on mitochondria biology. We found that Lin and Rif did not target mitochondrial translational machinery, as the levels of the mitochondrial-encoded NAD9 proteins were comparable or even higher in treated seedlings *versus* the control sample (Figure 2 (a) and Supplementary Figure 3), which confirms the findings of the previous study (Mulo *et al*., 2003). On the other hand, a reduction of around 40% of the NAD9 protein levels was observed for the Spec treatment (Figure 2 (a)), as we have previously documented (Sajib *et al*., 2023a). However, the accumulation of mitochondrial transcripts is not decreased by any of the treatments (Figure 2 (b)) but is slightly augmented for the Spec treatment, likely as compensation for the decrease in the mitochondrial protein levels. Consistent with the effect of Spec on mitochondrial translation, the accumulation of AOX1A transcript/AOX proteins and other nuclear marker transcripts for mitochondrial stress was induced only upon treatment with Spec but not with Lin (Figures 2 (a), (b) and Supplementary Figure 4). The induction of mitochondrial stress markers in Spec-treated seedlings was comparable or even higher for *Aox1a* transcripts than in the mutant *atphb3,* which is impaired in mitochondrial function and morphology (Van Aken *et al*., 2016). The usage of genetically encoded biosensors for monitoring changes in H_2_O_2_ and thiol redox balance (Nietzel *et al*., 2019) also indicates that treatment with Spec at high concentration or Rif but not with Lin induces the production of mitochondrial-localized ROS (Figure 2 (d) and Supplementary Figure 6). We conclude from the data that the antioxidant system in seedlings maintains the redox status only when seedlings are treated with a lower Spec dose. In the presence of higher Spec concentrations, the equilibrium between the production and scavenging of ROS may be perturbed, which can cause sudden increases in intracellular ROS levels (Huang *et al*., 2019).

Furthermore, metabolomics profiling was conducted on WT Arabidopsis seedlings treated with antibiotics to identify metabolites linked to PGE limitation and seedling growth (Figure 2 (c), Supplementary Figure 5). Analysis via GC-MS showed no alterations in TCA and glycolytic metabolites in Lin-treated seedlings. In contrast, Spec-treated seedlings exhibited an accumulation in steady-state levels, suggesting that Spec treatment rather than Lin alters mitochondrial respiration metabolism. Previous studies have demonstrated that in mitochondrial mutants and under conditions of mitochondrial metabolism perturbation, amino acids can accumulate, likely due to cellular energy deficits (Van Aken *et al*., 2016). In this study, we found a significant alteration in amino acid metabolism with Spec treatment and no changes in amino acid profiles with Lin treatment. The observed shift in amino acid metabolism with Spec suggests deficiency in ATP production, which is not compensated by photosynthesis in dark growth conditions, leading to impaired synthesis of aspartate (ASP), asparagine (ASN), and glutamine (GLN) (Supplementary Figure 5). On the other hand, the accumulation of pyruvate-derived branched amino acids in Spec-treated seedlings can be interpreted as the result of the accumulation of pyruvate due to TCA cycle limitation. These findings further support the notion that Spec affects mitochondrial energy supply, unlike Lin.

Alternatively, a decrease in ASP-derived ASN levels could be linked to plastid malfunction in Lin/Spec-treated seedlings, directing ASP flux to the synthesis of ILE. Finally, among shikimate-derived amino acids, only the anthranilate branch of the pathway seems to be upregulated in Spec treatment, as shown by an increase in TRP levels, which could impact auxin synthesis involved in apical hook formation.

Previous studies have shown that mitochondrial dysfunction mutants (e.g., *atphb3*, *rpoTmp*) exhibit altered levels of fatty acids and lactic acid (Van Aken *et al*., 2016). We also observed the accumulation of various fatty acids, e.g., palmitic acid and myristic acid, in Spec-treated seedlings but not in those treated with Lin (Suppl. Figure 5 (c)). Notably, lactic acid shows a tendency to accumulate in Spec-treated seedlings while not detected in Lin one (Suppl. Figure 5 (b)), indicating a shift to fermentation, a classical indicator of impaired mitochondrial respiratory function. All these observations highlight the impact of antibiotic treatments on metabolic pathways: the metabolic profile in Spec-treated seedlings signs a dual plastid/mitochondrial alteration with mostly mitochondrial alteration; meanwhile, in Lin treatment, the metabolic profile is almost unchanged. These data also indicate that plastids do not play a significant role in the cellular metabolism of etiolated seedlings, though they control skotomorphogenesis. We suppose that the amino acid biosynthetic pathway, that is mostly composed of nuclear-encoded enzymes, is untouched even if plastid gene expression is impaired by lincomycin. In addition, these enzymes are stromal and therefore not likely to be impacted by the alteration of the etioplast structure in Lin-treated seedlings. These data exclude an inhibitory action of Lin on mitochondrial translation and metabolism and show that Lin-induced plastid dysfunction is not accompanied by mitochondrial stress and/or mitochondrial ROS production. On the other hand, Spec treatment clearly impacts mitochondrial translation, metabolism, and ROS production. We confirm that seedling treatment with Lin is the most pertinent method for investigating the direct role of plastids on skotomorphogenesis.

We found that plastid translation inhibition by Lin induces an exaggeration of the apical hook bending that is comparable to the alteration induced by Spec (Figures 1(d) and (e)). When the GUN1 plastid retrograde pathway was genetically disrupted, most of the Lin-treated seedlings were untwisted (Figure 3 (b) and (c)). In addition, Lin treatment did not alter the expression of PhANGs in mutant *gun1* seedlings (Figure 3 (a)). These data indicate that the Lin-induced skotomorphogenic and nuclear expression reprogramming is GUN1 dependent. In the case of Spec treatment, which induces both plastidial and mitochondrial stress, mutant *gun1* seedlings still respond at the developmental and nuclear expression levels, even if in a weaker manner than the WT. These data suggest the involvement of other signaling pathways in addition to the GUN1-dependent signalization in the case of Spec. A recent study indicates that the ANAC017OE2 can maintain the expression of PhANGs upon Lin treatment (Zhu *et al*., 2024). Therefore, it will be interesting to explore whether the ANAC017-dependent retrograde pathway plays a role in regulating the expression of PhANGs in response to Spec treatment. Previously, in Rif-treated seedlings, we described the induction of marker transcripts for mitochondrial stress and the capacity of mitochondrial AOX-dependent respiration (Sajib *et al*., 2023a). In addition, the twisting phenotype was dependent on the activity of the mitochondrial AOX1A enzyme in both Rif and Spec-treated seedlings. All these data are indications of the involvement of mitochondrial activity in the response to PGE limitation by Rif and Spec. The expression analysis could not reveal any alteration of the PhANGs expression upon Rif treatment, and the developmental response was unchanged in mutant *gun1* seedlings. We propose that whenever plastidial dysfunction is accompanied by mitochondrial stress, the retrograde plastid-specific pathways, such as GUN1, are either hidden or under-used, leaving space for other signaling mechanisms, eventually related to mitochondria. Recently, it was documented that an increase in mitochondrial ROS is involved in the activation of nuclear reprogramming in mitochondrial stress circumstances (Khan *et al*., 2024). However, the mechanisms operating downstream to the ROS accumulation are still undefined.

It was shown that the ethylene signaling pathway is involved in regulating the bending of the apical hook in etiolated seedlings in darkness (Guzman and Ecker, 1990) and in underground conditions (Zhong *et al*., 2014). However, previously, we have documented that overbending of the apical hook was still stimulated by Rif, even in mutant *ein2* seedlings (Sajib *et al*., 2023a). In addition, ethylene-responsive genes were not induced, and ethylene levels did not accumulate. In our current study, we could also show that *ein2* mutants were still responding at the apical hook bending level to Spec and Lin treatments (Figure 3 (d) and (e)), indicating that the EIN2-dependent pathway is not involved. However, the response to Lin was shown to act independently of EIN2 and downstream to EIN3/EIL1 in light-grown seedlings (Gommers *et al*., 2021). Alternatively, the ethylene signaling pathway can also use not canonical components that can be parallel to the EIN2 core pathway, such as AHP (Arabidopsis Histidine-containing Phosphotransferase protein) and ARR (Arabidopsis Response Regulator) (Binder, 2020). Further research will be required to test these hypotheses in the context of Spec and Lin.

In conclusion, a developmental response at the level of the apical hook bending was observed whenever organellar functions were limited in dark-growing seedlings. The use of KCN treatment targeting the mitochondrial electron transfer chain and mutations affecting mitochondrial biology (as in *rug3* and *atpHB3*) could restrict the developmental response to mitochondrial dysfunction (Merendino *et al*., 2020). On the other hand, whenever plastid gene expression was limited using Rif or Spec treatments, marks of mitochondrial stress were also detected as induction of stress response gene expression levels, production of mitochondrial ROS, and alteration of mitochondrial metabolites, making it difficult to distinguish a plastid-specific response from a mitochondrial one (Figure 4). Here, we show that Lin treatment induces changes in skotomorphogenesis by inhibiting plastid translation, specifically, without affecting mitochondria biology, which underscores the function of plastids in controlling early development, even in the absence of photosynthetic activity.

**Figure 4.**
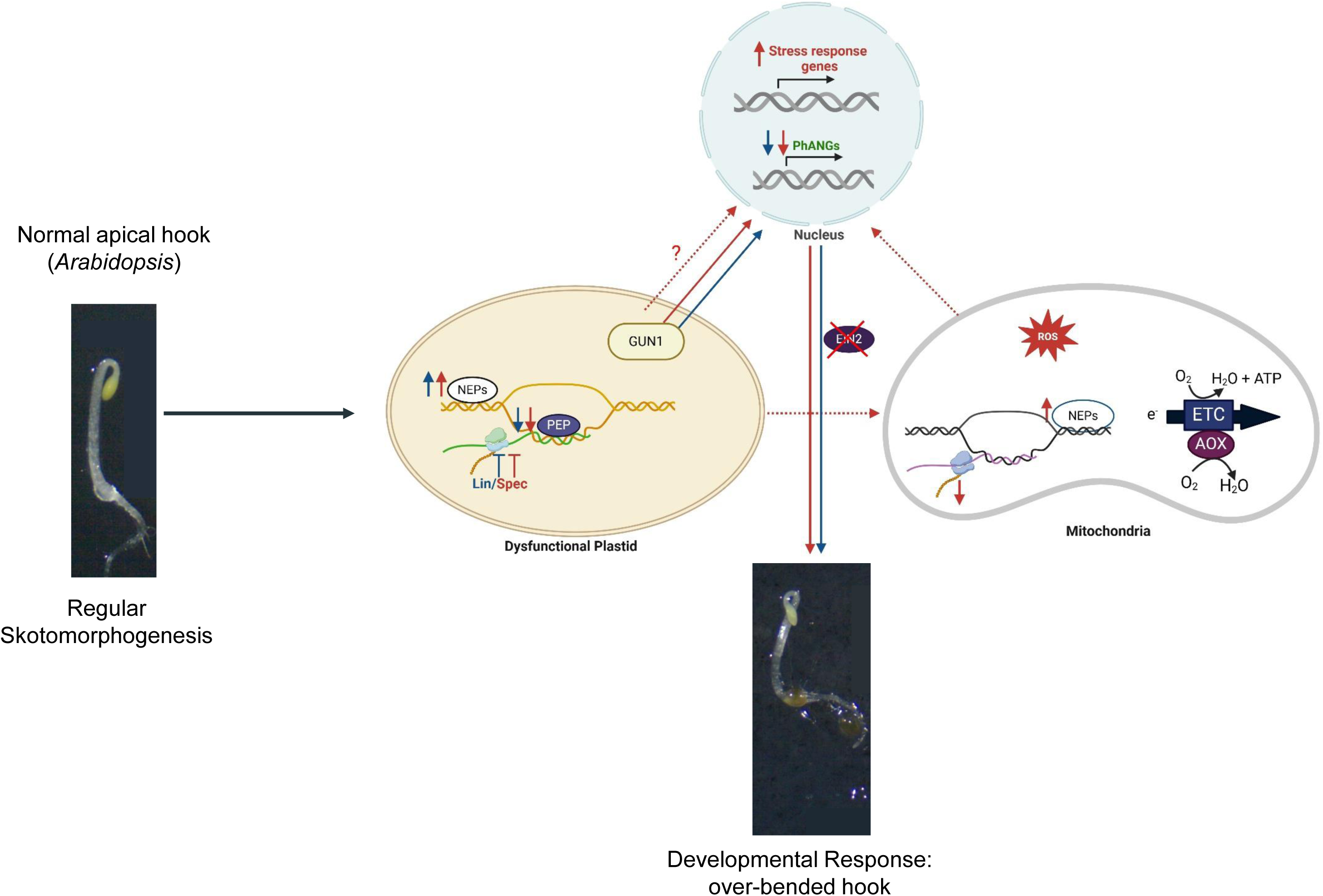
Scheme of the proposed functional link between plastid translation inhibition by Lin/Spec and skotomorphogenic reprogramming. In etiolated WT seedlings that are treated with Lin (blue lines) or Spec (red lines), the translation of plastid transcripts and, subsequently, the transcription of the PEP-dependent plastid genes are inhibited. Consequently, the NEP-dependent expression of plastid genes is induced as compensation, and the levels of Photosynthesis-Associated Nuclear Genes (PhANGs, in green) are down-regulated by the GUN1-mediated plastidial retrograde pathway. Response to Spec treatment is also mediated by unknown GUN1-independent signalization (red dotted arrows). Spec, but not Lin, alters mitochondrial genome expression, by inhibiting mitochondrial translation and up-regulating mitochondrial transcripts, and metabolism; at the same time, Spec induces the expression of nuclear markers for mitochondrial stress and metabolism (red dotted line). Consequently, the mitochondrial redox status in the cell is altered, and mitochondrial ROS is produced. Whether the impact of Spec on mitochondrial biology is direct or indirect is a matter of further exploration. The ultimate result of the plastid translation inhibition by Lin and Spec consists in skotomorphogenic reprogramming (twisting phenotype: apical hook over-bending) that is independent of the EIN2-mediated pathway.

## Material and Methods

### Plant material

*Arabidopsis thaliana* wild-type (WT) (ecotype Columbia) and mutant lines-*atphb3* (SALK_020707), *gun1-201* (SAIL_290_D09^24^), *ein2-1* [CS3071; (Guzman and Ecker, 1990)] and mitochondrial roGFP2-Orp1 line from (Nietzel *et al*., 2019) were used in this study. Sterilized seeds are placed on Murashige and Skoog agar plates containing 1% sucrose and 0.08% charcoal (a black powder that enhances the background picture of etiolated plants, SIGMA-ALDRICH). Lin (Sigma-Aldrich 62143) and Spec (Sigma-Aldrich S4014) were added to MS agar plates, with a final concentration of 1 mM and 0.75 mM, while MS agar plates were used as a mock control. To identify the mitochondrial redox status, 1.5 mM Spec was used (Figure 2 (d)). Rif (Sigma-Aldrich R3501) was added to a final concentration of 0.12 mM (100 mg/ml) and 0.24 mM (200 mg/ml), while MS agar plates containing DMSO were used as a mock control. After that, plates containing the sown seeds are stratified for three days at 4°C in the dark, exposed for six hours to light (100 µEm^-2^ s^-1^ white light) at 20°C, and then grown in the dark at 20°C. After three days, the seedlings are harvested under green light for molecular analyses and phenotyping or the following experiments.

### Quantification and statistical analysis of phenotypic parameters

The phenotypic characteristics of the seedlings were analyzed as previously described (Merendino *et al*., 2020). Briefly, digital images of individual plants were taken with a stereo microscope (ZEISS Stemi 305, Switzerland) using the ZEN 2.3 software. Next, we used ImageJ to determine the angles developed in the apical hook between the hypocotyl and cotyledons. Any hook angle larger than 180° was considered as twisted. All experimental data were analyzed using a linear model that accounted for both the biological effects and the potential influence of replicate effects. The Shapiro-Wilk normality test was applied to the data to assess the normality of the residuals. In cases where the Shapiro-Wilk test indicated non-normality, the observations were split into two subpopulations above or below an angle of 180° within each replicate. Finally, for each replicate, Pearson’s Chi-squared test with Yates’ continuity correction was applied to evaluate the significant differences in the proportions of the subpopulations between the two conditions. All statistical analyses were conducted using R version 4.1.2.

### Transmission Electron Microscopy (TEM)

Samples for transmission electron microscopy (TEM) were fixed for 2 hours in 2.5% glutaraldehyde in 50 mM cacodylate buffer (pH 7.4). After fixation, they were post-fixed in 2% OsO4 at 4°C overnight and then dehydrated using a graded series of acetone. Dehydrated samples were infiltrated with acetone: resin mixtures (3:1, 1:1, 1:3) and finally embedded in a pure resin medium (Agar 100 Resin Kit). Polymerized blocks containing the samples were cut into 90 nm-thick sections using the Leica UCT ultramicrotome. The microscopy was performed in the Laboratory of Electron Microscopy of the Nencki Institute, supported by the project financed by the Minister of Education and Science based on contract No 2022/WK/05 (Polish Euro-BioImaging Node “Advanced Light Microscopy Node Poland”) using the JEM 1400 (JEOL) equipped with a Morada G2 CCD camera (EMSIS GmbH). The ultrastructural characteristics of the PLB were calculated using ImageJ software (Abramoff et al., 2004). Cross-sectional areas of PLB and starch grains were measured from 2D images of whole etioplasts by manually tracing the respective structures visible in the micrographs. Additionally, plastoglobule diameters were determined through manual measurements at the midpoint of visible plastoglobule structures. The average number of starch granules per etioplast cross-section was established by manual counting based on images of whole etioplasts.

### Protochlorophyllide (Pchlide) quantification

The level of protochlorophyllide (Pchlide) was quantified using the Shimadzu RF-5301PC spectrofluorometer with the protocol described earlier (Zhong et al., 2014). In brief, 50 dark-grown etiolated seedlings were collected under dim green, safe light and ground in liquid nitrogen, followed by vortexing in 1 mL of extraction buffer consisting of 90% (vol/vol) acetone with 0.1% ammonia (NH_3_). After centrifugation, the fluorescence emission spectra of the supernatant were analyzed at room temperature. The excitation wavelength was 440 nm, and the spectra of emission from 610 to 750 nm were recorded with a 1 nm bandwidth. The relative abundance of Pchlide was calculated by comparing the highest fluorescence value of each condition with the controls. All Pchlide measurements were conducted independently in 3 biological triplicates. Statistical significance was determined using one-way ANOVA with post hoc Tukey test at p < 0.05.

### RT-qPCR analysis of RNA levels

Total RNAs were isolated from the grounded whole seedling using the NucleoSpin RNA Plus Mini kit (Macherey-Nagel, Germany) using the manufacturer’s protocol. In brief, after homogenizing and lysing the grounded plant material, genomic DNA was removed by filtrating. The lysate was then used to bind the RNA-binding column, followed by washing with the buffer. The RNA was then eluted into RNase-free H_2_O. 500 ng of RNA was reverse transcribed using the Maxima H Minus Reverse Transcriptase (Thermo Fisher Scientific Baltics UAB, Lithuania). Random primers were used in order to detect nuclear, plastidial, and mitochondrial transcripts. Using the LightCycler® 480 SYBR Green I Master mix (Roche Life Science, Germany) and forward and reverse primers (0.5 µM; Suppl. Table 1), the qPCR reaction was performed in a Biorad CFX384^TM^ Real-Time System PCR machine (Biorad, USA). The data was then analyzed using the CFX Manager Software. PP2A was utilized as a reference gene for normalizing transcript expression levels. The mean values of the biological replicates were shown, and standard deviation are represented as error bar.

### Multiwell plate reader-based fluorimetry

For the plate reader assays, pools of 3-day-old mito-roGFP2-Orp1 seedlings (10 seedlings per pool, 60 seedlings per sample) were placed in 100 µL plate reader buffer (10 mM MES, 10 mM MgCl_2_, 10 mM CaCl_2_, 5 mM KCl, pH 5.8) in a 96-well plate (black, flat bottom, Greiner bio-one, Frickenhausen, Germany) (Nietzel *et al*., 2019). Fluorescence measurements were conducted using a CLARIOstar plate reader (BMG Labtech, Germany) set to 25°C. Measurements were performed with spiral averaging mode, comprising 54 excitation light flashes along a 6 mm radius circle without shaking. The seedlings were excited sequentially with monochromatic light at 400 ± 10 nm and 482 ± 16 nm, followed by the emission detection at 520 ± 10 nm. After 3h of measurements, the plate reader medium was supplemented with either 50 mM H_2_O_2_ or 100 mM DTT to achieve complete oxidation or reduction of the sensor, respectively. The plate was reinserted into the plate reader to resume measurement. To avoid the effects of light, all the experiments were performed under safe (green) light. Fluorescence intensity was used to determine the ratiometric measurement of the mito-roGFP2-Orp1 sensor. Background corrections were performed at each time point by subtracting the autofluorescence of wild-type (WT) seedlings (included on the same plate) from the acquired sensor fluorescence data. Following that, quantification of the degree of oxidation (OxD) of the sensor was performed from the following equation described earlier (Schwarzländer *et al*., 2008):

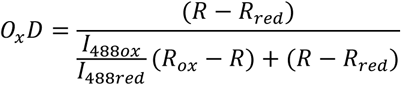

Where *R* represents the excitation ratio at 400/482 nm, *R_red_* is the ratio of the fully reduced form subsequent to perfusion with 100 mM DTT; *R_ox_* indicates the ratio of the fully oxidized form after perfusion with 50 mM H_2_O_2_ and *I_482ox_* and *I_482red_* are the intensities at 482 nm for the fully oxidized and fully reduced forms, respectively. Statistical differences between control and Lin/Spec/Rif treated seedlings were determined using a two-tailed unpaired Student’s t-test.

### Metabolic analyses

Metabolic profiling was obtained using GC-MS. Frozen (−20°C) water:acetonitrile: isopropanol (2:3:3) containing 4 µg mL^-1^ of Ribitol was used for the resuspension of ground fresh samples (50 mg FW). The mixture was then shaken for 10 minutes at 4°C, centrifuged, and 100µl of the supernatant (SN) was dried for 4 hours at 35°C in a Speed-Vac before being stored at −80°C. All GC-MS analysis procedures were carried out following the instructions described earlier (Fiehn, 2006; Fiehn et al., 2008). The content of the metabolites was expressed in arbitrary units (Semi-quantitative determination). Amino acids quantification was performed using OPA-HPLC profiling according to the protocol described earlier (Sajib et al., 2023). The content of the metabolites and amino acids was normalized with the control sample of each replication. Each experiment was independently replicated four times. The statistical significance of the metabolites was identified using a one-way ANOVA test with a post hoc Tukey test with a significance level set at P < 0.05.

### Western blot analysis

For the extraction of total protein, 100 mg of ground plant material was resuspended in 100 μl of protein lysis buffer (50 mM Tris pH 8.0; 2% SDS; 10 mM EDTA; protease inhibitors (cOmplete™, Mini, EDTA-free Protease Inhibitor Cocktail, Roche (04693159001), Sigma-Aldrich)) and then incubated at room temperature for 30 min. Then, the samples were centrifuged for 30 min at 12000 × g at 4°C, followed by collection of the supernatant (SN). Then, the amount of protein in the sample was quantified by BCA assay using the Bicinchoninic acid solution (Sigma-Aldrich B9643) and CuSO₄.5H₂O. The samples were then denatured for 5 min at 95 °C in 4X reducing electrophoresis sample buffer composed of 200 mM Tris pH 6.8, 5% mercaptoethanol, 4% SDS, 0.2% Bromophenol Blue, and 20 % glycerol. 50 μg of plant protein extracts were separated using SDS-PAGE (12% acrylamide), electroblotted onto a milk-saturated nylon membrane, and immunodetected using specific polyclonal rabbit antisera against the plant mitochondrial-encoded NAD9 (Lamattina et al., 1993), the plastidial-encoded RBCL (Agrisera, AS03 037), plant nuclear-encoded mitochondrial alternative oxidase 1 and 2 (AOX1/2; Agrisera, AS04 054), and the nuclear-encoded vacuolar V-ATPase (Agrisera, AS07 213). Finally, the blots were developed with Chemiluminescent Peroxidase Substrate-3 (CPS3100, Sigma-Aldrich). A CCD imager (Chemidoc MP, Bio-Rad) and the Image Lab software (Bio-Rad, Hercules, USA) were used to capture images of the blots.

## Supporting information

primers and GCMS data

## Acknowledgments

This work was supported by the LabEx Saclay Plant Sciences-SPS (ANR-10-LABX-0040-SPS) to IPS2 and LM (ANR-11-IDEX-0003-02); EUGLOH-UPSaclay Research program (ANR-19-GURE-0006). TEM and Pchlide spectroscopy studies were funded by the National Science Centre, Poland, under the OPUS call in the Weave program (2022/47/I/NZ3/00498). We thank Robert Blanvillain, Michael Hodges, Emmanuelle Issakidis-Bourguet, and Olivier Van Aken for helpful discussions. SAS was supported by a fellowship from the Ministère de l’Enseignement Supérieur, de la Recherche et de l’Innovation (MESRI) of the French Government (Doctoral School of Plant Sciences [SEVE], Université Paris-Saclay, France) for his PhD. The scheme in Figure 4 was created in BioRender (Sajib, S. (2024) BioRender.com/v19d372).

## Author contribution

LM and BG designed research; SAS, MB, PB, CM, RC, and LK performed experiments; ED performed statistical analyses; SAS, MB, PB, CM, MS, LK, BG, and LM analyzed data; SAS and LM wrote the manuscript with the help of all co-authors.

---

**Suppl. Figure 1.**
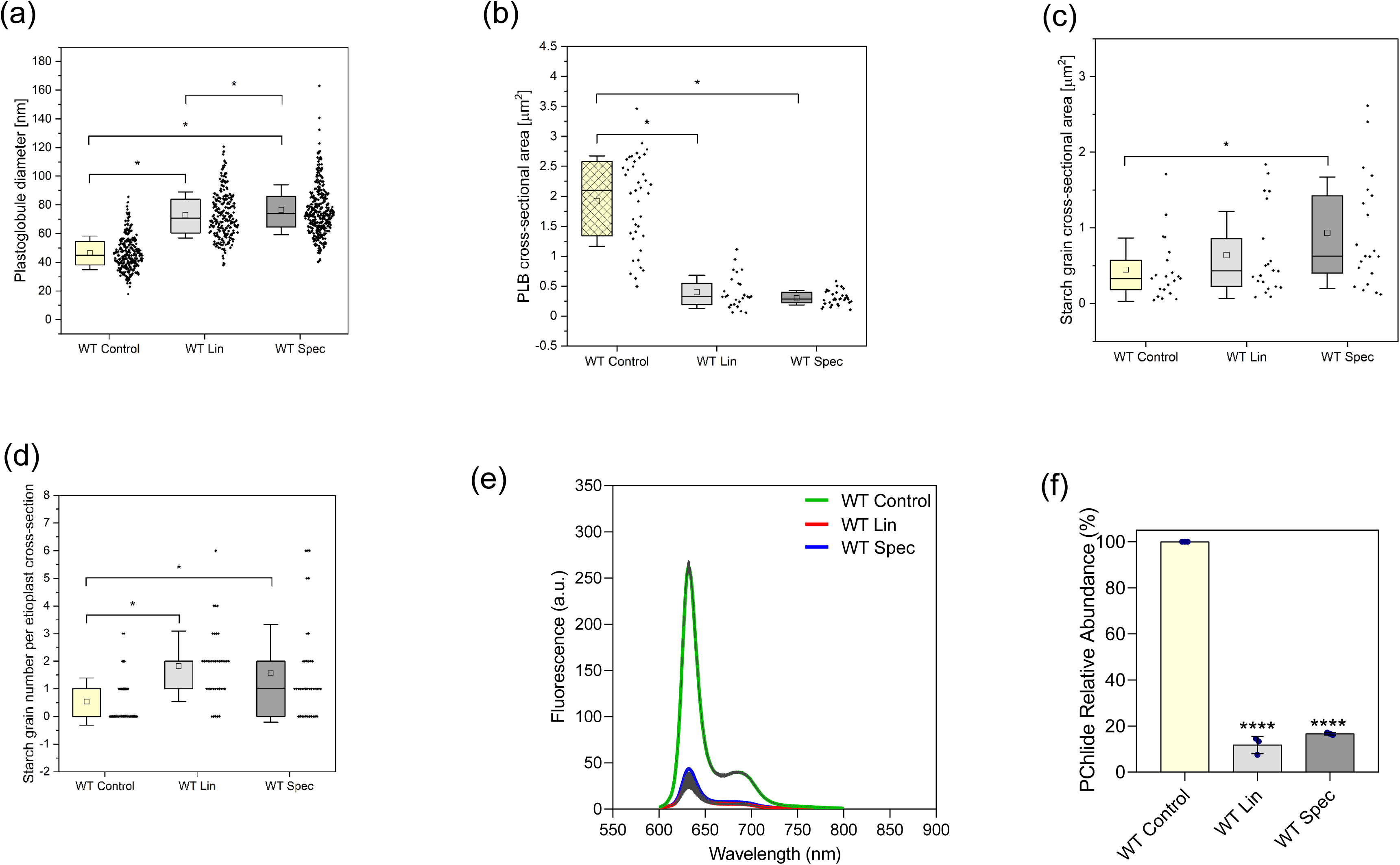
Measurements of etioplast structural parameters: (a) Plastoglobule diameters (n = 251-364), (b) PLB cross-sectional area (n = 26-39), (c) Starch grain cross-sectional area (n = 21), (d) Number of starch grains per etioplast cross-sectional area (n = 39). All measurements were performed on images obtained from three biological replicates. (e) Protochlorophyllide (Pchlide) fluorescence at 634 nm was measured in two biological replicates of 3-day-old dark-grown seedlings, with results presented as mean ± SD; n = 3 (50 seedlings per replicate). (f) The relative abundance of Pchlide was calculated by comparing the highest fluorescence values of each type of treatment with the controls. Statistical significance was determined using one-way ANOVA with post hoc Tukey test (*P < 0.05, ****P < 0.0001).

**Suppl. Figure 2.**
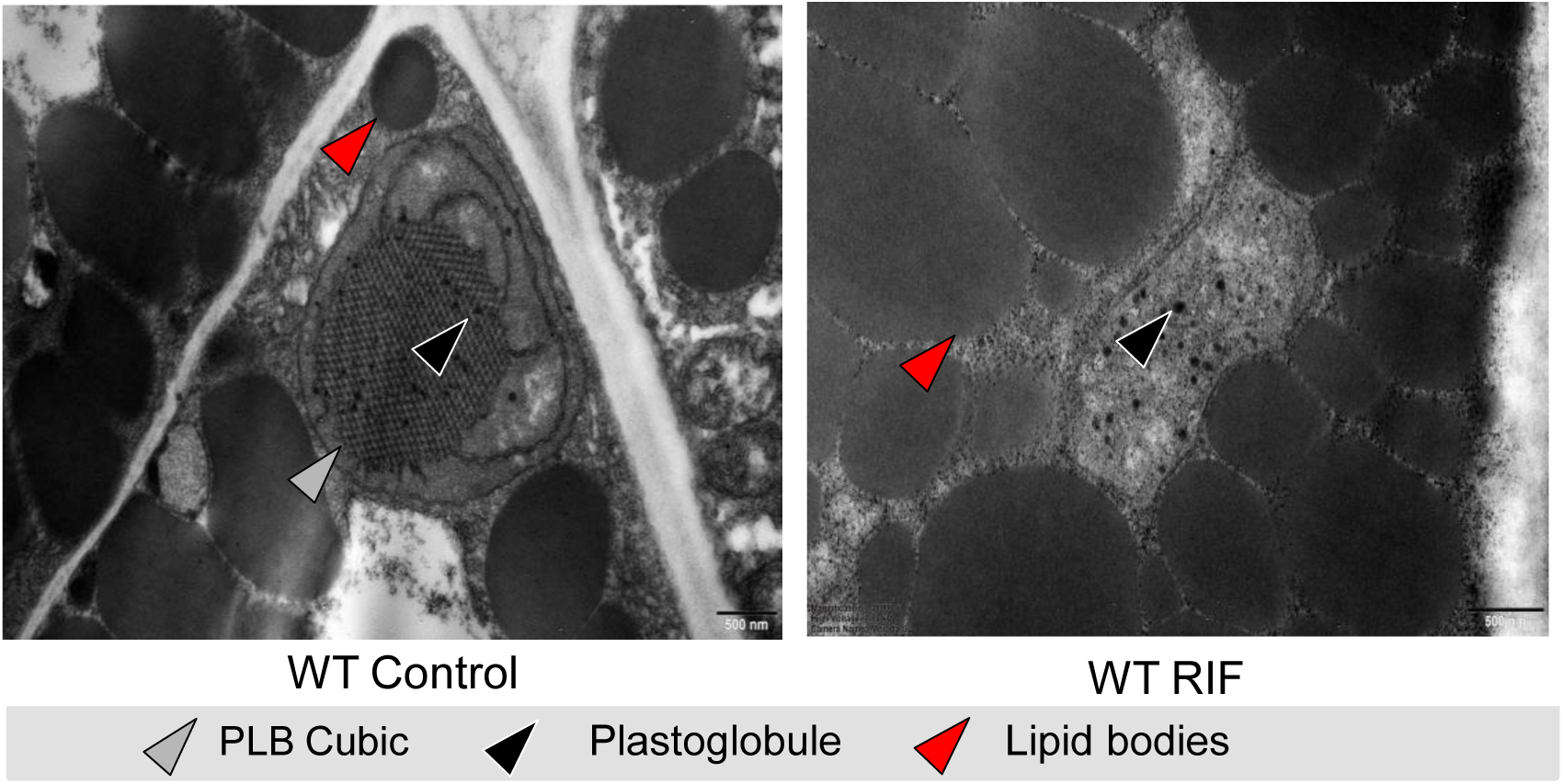
Ultrastructure of etioplasts in cotyledon regions of 3-day-old etiolated Arabidopsis WT seedlings grown in the presence of DMSO (control) or 200 µg/ml Rif. Grey arrowhead with black outline indicates the cubic prolamellar body (PLB), black arrowhead with white border indicates plastoglobules and red arrowhead with black border indicates lipid bodies. The scale bar is 500 nm.

**Suppl. Figure 3.**
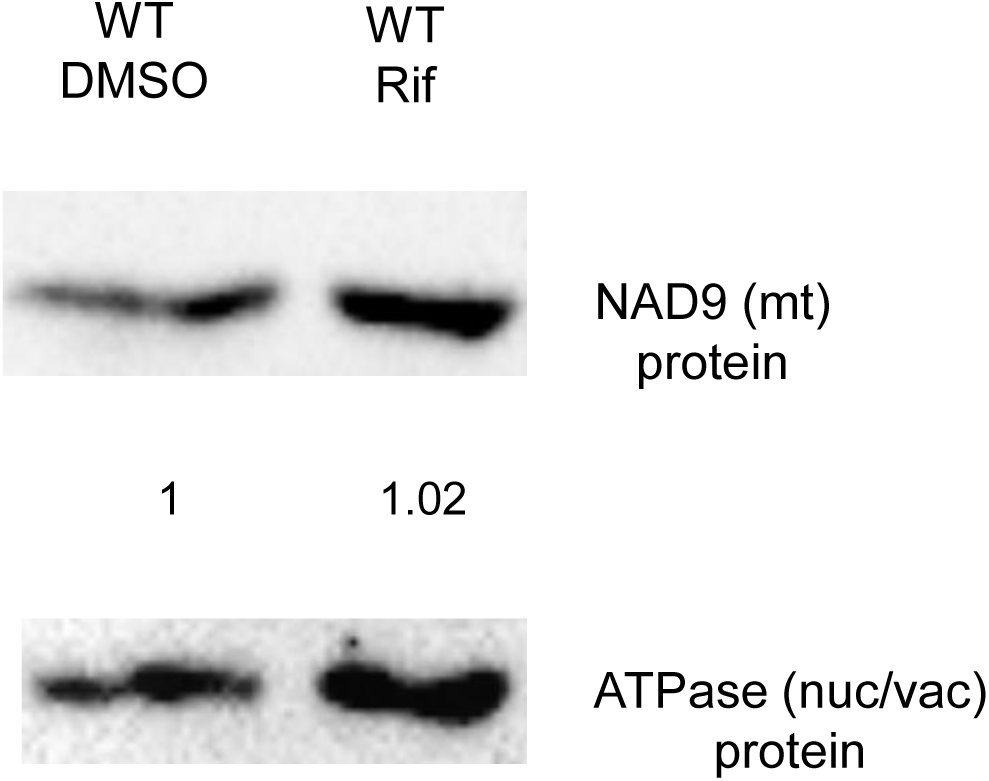
Analysis of mitochondrial-encoded NAD9 protein content: Protein extracts (30 µg) from etiolated WT seedlings grown in the presence of DMSO (control) or 200 µg/ml Rif were separated by SDS-PAGE and immunoblotted with specific antisera against mitochondrial-encoded NAD9 and the vacuolar protein ATPase (nucleus-encoded) as a loading control. Quantification of the NAD9 signals was performed using Image Lab software with ATPase for normalization; values are reported below the NAD9 protein signals.

**Suppl. Figure 4.**
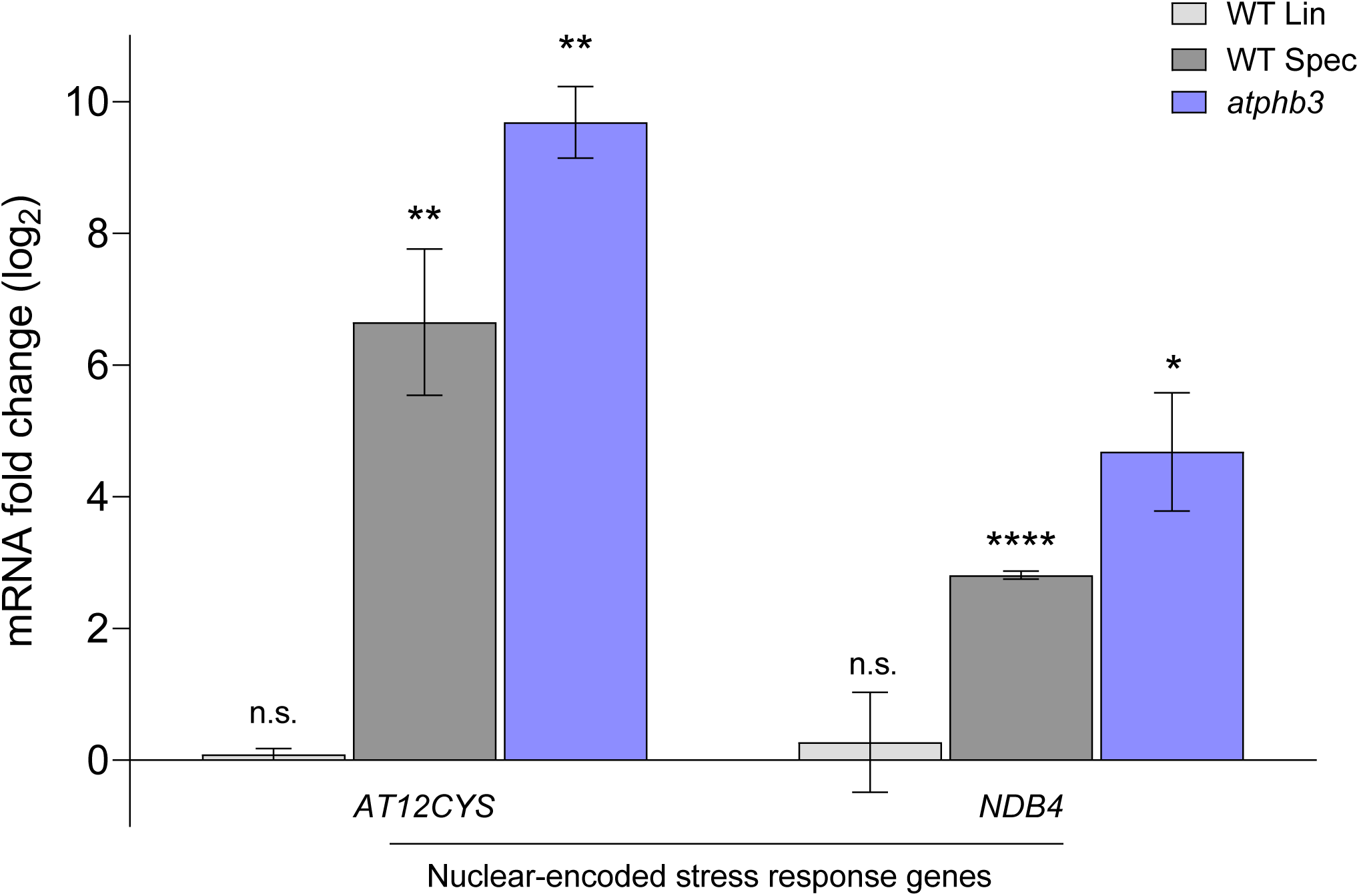
Transcript levels in 3-day-old etiolated Arabidopsis seedlings: Log_2_(FC) values of expression levels of nucleus-encoded stress marker genes were determined by reverse transcription-quantitative PCR in Lin/Spec-treated WT seedlings and untreated mitochondrial mutant *atphb3* versus untreated WT seedlings. Expression levels were normalized to the mean expression of the nuclear reference gene *PP2A*. The mean values of three biological replicates are plotted, with error bars representing standard deviation. The statistical significance of differences was determined using a t-test (*P < 0.05; **P < 0.01, ****P < 0.0001, n.s. = not significant, n.d.= not detected).

**Suppl. Figure 5.**
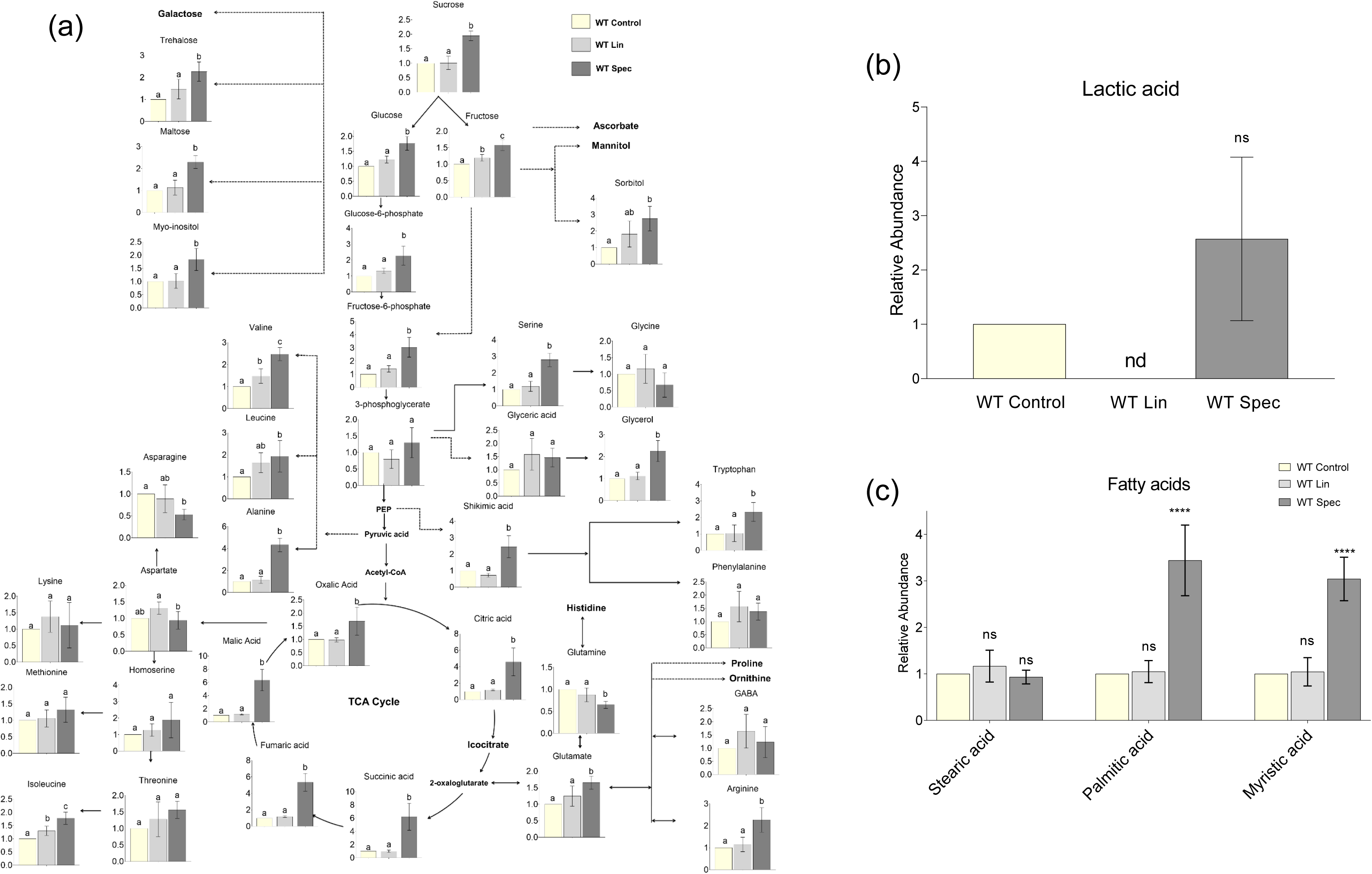
Impact of plastid translation inhibition on metabolism. (a) Schematic representation of the relative abundance of metabolites (determined by GC-MS) and amino acid levels (determined by OPA-HPLC). Statistical significance was determined using one-way ANOVA with post hoc Tukey test. Different letters indicate significant differences among the samples (P < 0.05). (b) Relative abundance of lactic acid and (c) fatty acids (determined by GC-MS) in untreated and Lin/Spec-treated WT seedlings (4 biological replicates). Error bars represent standard deviation. Statistical significance was determined using one-way ANOVA with post hoc Tukey test. (****P < 0.0001, ns = not significant, nd= not detected).

**Suppl. Figure 6.**
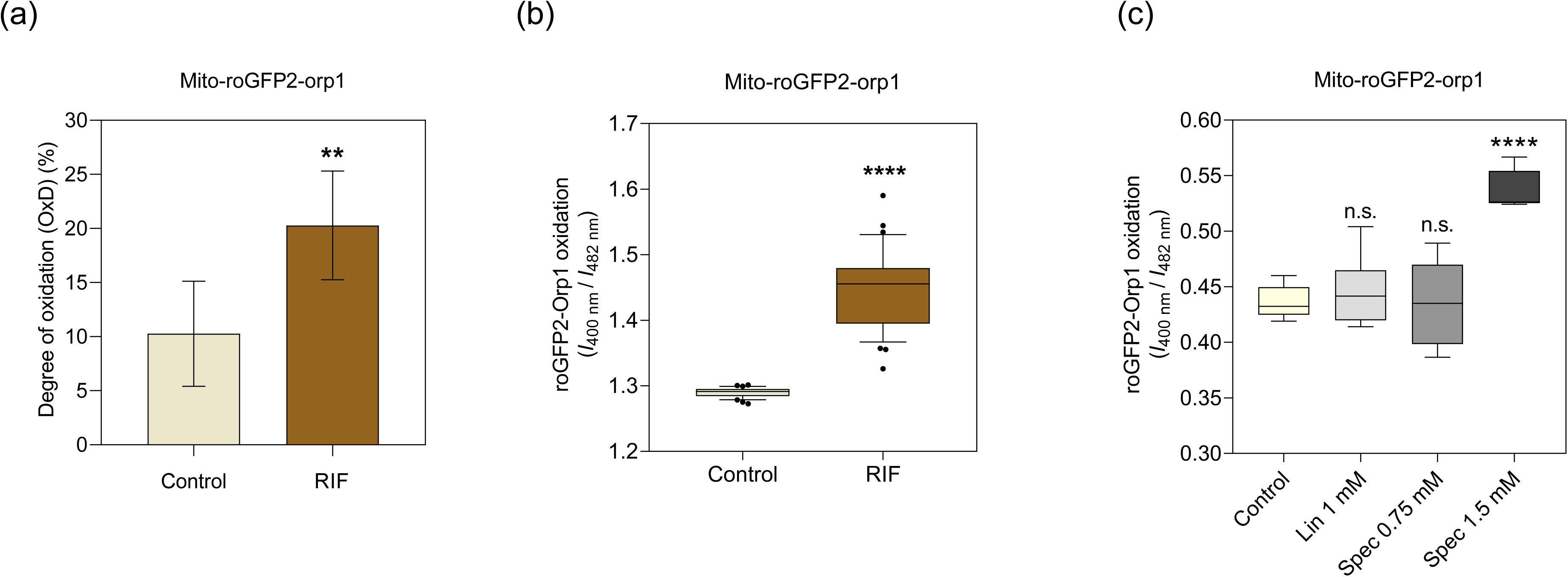
(a) Measurements of H_2_O_2_ and thiol redox status in the mitochondrial matrix using the mito-roGFP2-Orp1 line in DMSO (control) and 400 µg/ml Rif treated conditions. The degree of oxidation was calculated from measured ratios after in situ calibration with 50 mM H_2_O_2_ and 100 mM DTT (n = 80; significantly different with **P < 0.01, two-tailed, unpaired Student’s t-test). (b) Sensor steady-state oxidation levels of the mito-roGFP2-Orp1 line in DMSO control and Rif-treated seedlings, and (c) in control, 1 mM Lin, and 0.75 mM Spec and 1.5 mM Spec treated conditions were monitored under physiological conditions and after calibration with 100 mM DTT or 50 mM H_2_O_2_. n = 80; boxplots show 10–90% percentiles; n.s.= not significant, significantly different with **P < 0.01, ****P < 0.0001, two-tailed, unpaired Student’s t-test.

**Suppl. Figure 7.**
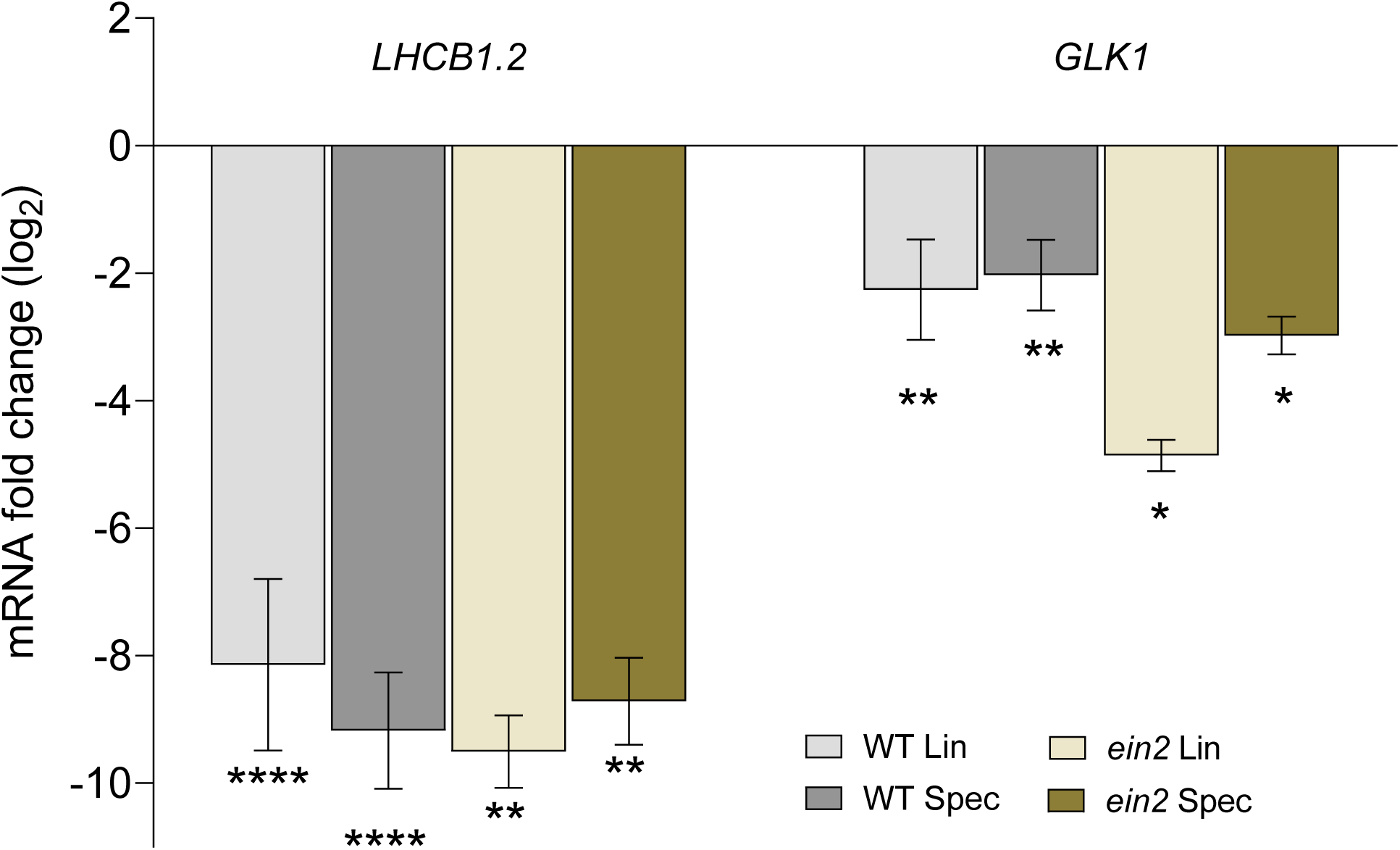
Log_2_ (FC) values of expression levels of Photosynthesis-Associated Nuclear Genes (PhANGs) in Lin/Spec-treated WT and mutant *ein2* seedlings versus corresponding untreated seedlings as determined by reverse transcription-quantitative PCR. The expression levels were normalized to the mean expression of PP2A, which was used as a reference gene. The mean values of three biological replicates are plotted. Error bars correspond to standard deviation. The t-test was performed to determine the statistical significance of differences (*P < 0.05, **P<0.01, ****P < 0.0001).

